# *Plasmodium falciparum* exploits CD44 as a co-receptor for erythrocyte invasion

**DOI:** 10.1101/2023.04.12.536503

**Authors:** Barbara Baro-Sastre, Chi Yong Kim, Carrie Lin, Angel K. Kongsomboonvech, Marilou Tetard, Nichole D. Salinas, Niraj H. Tolia, Elizabeth S. Egan

## Abstract

The malaria parasite *Plasmodium falciparum* invades and replicates asexually within human erythrocytes. CD44 expressed on erythrocytes was previously identified as an important host factor for *P. falciparum* infection through a forward genetic screen, but little is known about its regulation or function in these cells, nor how it may be utilized by the parasite. We found that CD44 can be efficiently deleted from primary human hematopoietic stem cells using CRISPR/Cas9 genome editing, and that the efficiency of ex-vivo erythropoiesis to enucleated cultured red blood cells (cRBCs) is not impacted by lack of CD44. However, the rate of *P. falciparum* invasion was substantially reduced in CD44-null cRBCs relative to isogenic wild-type (WT) control cells, validating CD44 as an important host factor for this parasite. We identified two *P. falciparum* invasion ligands as binding partners for CD44, Erythrocyte Binding Antigen-175 (EBA-175) and EBA-140, and demonstrated that their ability to bind to human erythrocytes relies primarily on their canonical receptors-glycophorin A and glycophorin C, respectively. We further show that EBA-175 induces phosphorylation of erythrocyte cytoskeletal proteins in a CD44-dependent manner. Our findings support a model where *P. falciparum* exploits CD44 as a co-receptor during invasion of human erythrocytes, stimulating CD44-dependent phosphorylation of host cytoskeletal proteins that alter host cell deformability and facilitate parasite entry.

## INTRODUCTION

Malaria due to the Apicomplexan parasite *Plasmodium falciparum* is a leading cause of morbidity and mortality in the developing world, responsible for ∼627,000 deaths and millions of cases per year ^1^. The clinical symptoms of malaria occur when the parasites invade and replicate asexually in erythrocytes (red blood cells; RBCs) in exponential cycles lasting ∼48 hours. Asexual replication is critical to disease pathogenesis and thus is a major focus of intervention efforts. Yet, emerging drug resistance and the lack of a strongly effective vaccine remain major obstacles to global malarial control. A deeper understanding of the host-pathogen interactions occurring during the blood stage of this complex parasite’s life cycle is needed to advance new therapeutic or preventive strategies ^2–5^.

*P. falciparum* follows a multi-step process for successful erythrocyte invasion, including apical reorientation, host cell deformation, and formation of a moving junction at the cellular interface ^6^. The driving force for internalization is powered by the parasite’s actino-myosin motor. Like other invasive parasites within the phylum Apicomplexa, *P. falciparum* merozoites harbor two major secretory organelles at their apical tip, the micronemes and the rhoptries, in which a variety of invasion proteins are localized. Secreted ligands from the erythrocyte binding-like (EBL) and reticulocyte binding protein homologs (PfRh) protein families bind to specific receptors on the erythrocyte, including glycophorin A, B, C and CR1 ^2–5^. EBA-175 and EBA-140 are *P. falciparum* EBL ligands that interact with RBCs by binding to glycophorin A (GYPA) and glycophorin C (GYPC), respectively ^2, 7–14^. Binding occurs through the RII region of each EBA protein, which consists of two Duffy binding-like (DBL) domains ^9, 13–15^. The EBL and Rh ligands are considered functionally redundant, and mediate apical reorientation and host cell deformation during attachment ^16^. Following apical re-orientation, the *P. falciparum* ligand PfRH5 binds to its receptor, basigin. This interaction that has been proposed to result in a pore in the RBC membrane and to be required for rhoptry discharge and formation of the moving junction ^17–19^.

Our understanding of the role of erythrocyte host factors during *P. falciparum* invasion is limited, in part because of the genetic intractability of these unique cells, which are enucleated. CD44 was recently discovered as a host factor for *P. falciparum* through a forward genetic shRNA screen in cultured red blood cells (cRBCs) derived *ex-vivo* from nucleated hematopoietic stem/progenitor cells (HSPCs) ^20^. Further evidence for a role of CD44 in invasion was obtained using erythroblasts derived from the erythroleukemia cell line JK-1, where CD44-null JK-1 cells were resistant to invasion by three different *P. falciparum* strains ^21^. These results suggest that human CD44 is required for *P. falciparum* invasion in a strain-transcendent manner, yet its precise function remains unknown.

Widely expressed on eukaryotic cells, CD44 exists in several isoforms, but RBCs only express the shortest, standard form (CD44s) ^22^. Its function in human RBCs is largely uncharacterized aside from encoding the Indian blood group antigens ^23^. In other cells, the principal ligand for CD44 is hyaluronic acid (HA), which binds to the link domain, activating cell signaling pathways that modulate cytoskeletal changes and influence processes such as cell migration, metastasis, and survival ^24–28^. CD44 can serve as a receptor for Group A *Streptococcus* on epithelial cells, interacting with the bacterium’s HA capsule to stimulate signaling involving Rac1, phosphorylation of cellular proteins, and cytoskeletal rearrangements permissive for invasion ^29^. Through its cytoplasmic domain, CD44 interacts with Band 4.1 superfamily proteins such as Ezrin, Radixin, and Moesin (ERM) ^30, 31^. These interactions have been shown to lead to the activation of c-MET through Ras-MAPK and Pi3K-AKT pathways ^32–34^, but which downstream effectors are activated by CD44 depend on the stimulus and cell context ^28^. CD44 can also function as a co-receptor to help activate receptor tyrosine kinases in cancer cells and during *Listeria* infection of epithelial cells ^34–36^, but in neither case are the mechanisms fully understood.

Here, we investigated the function of erythrocyte CD44 during *P. falciparum* invasion using a combination of genetic, cell biological, and biochemical approaches. Our findings support a model where *P. falciparum* exploits CD44 as a co-receptor to facilitate its invasion into erythrocytes.

## METHODS

### *P. falciparum* culture

*P. falciparum* strain 3D7 is a laboratory-adapted strain obtained from the Walter and Eliza Hall Institute (Melbourne, Australia) and was routinely cultured in human erythrocytes isolated from de-identified peripheral blood erythrocytes (Stanford Blood Center) at 2% hematocrit in complete RPMI-1640 supplemented with 25 mM HEPES, 50 mg/L hypoxanthine, 2.42 mM sodium bicarbonate and 0.5% Albumax (Invitrogen) at 37° C in 5% CO_2_ and 1% O_2_.

### Antibodies

Antibodies used in this work include: anti-CD44 BRIC 222 mouse monoclonal (IBGRL), anti-CD44 rabbit polyclonal (Invitrogen PA5-21419), anti-GYPA-FITC clone 2B7 mouse monoclonal (Stem Cell Tech 60152FI), anti-GYPC-FITC mouse monoclonal (IBGRL), anti-6x-His mouse monoclonal (Invitrogen 37-2900), anti-CD99 mouse monoclonal (Invitrogen MA5-12954), anti-*P. falciparum* RAP1/2 mouse monoclonal (209.3; gift from Anthony Holder), R1454 anti-EBA-175 (3D7) rabbit anti-serum, R228 anti-EBA-140 rabbit anti-serum (gifts from Alan Cowman), anti-CD233 cytoplasmic domain BRIC 170 (IBGRL), goat anti-mouse IgG Alexa Fluor 488 (Invitrogen A-11001), goat anti-rabbit IgG Alexa Fluor 555 (Invitrogen A21428), goat anti-mouse IgG-FITC (Santa Cruz sc-2078) and goat anti-rabbit-HRP (Santa Cruz sc-2004).

### Ex-vivo erythropoiesis of primary human CD34^+^ HSPCs

Generation of cRBCs was performed essentially as previously described ^37^. Briefly, bone marrow primary human CD34^+^ HSPCs (Stem Cell Technologies; AllCells) were cultured in PIMDM composed of Iscove Basal Medium (Sigma) with 4 mM L-Glutamine, 330 µg/ml holo-transferrin, 10 µg/ml of recombinant human insulin, 2 IU/ml heparin, 10^-6^ M hydrocortisone, and supplemented with 100 ng/ml SCF, 5 ng/ml IL-3, 3 IU/ml Epo and 5% human plasma (Octaplas) at 37° C in 5% CO_2_. On day 2, the cells were nucleofected with ribonucleoprotein (RNP) complexes for CRISPR/Cas9-based genetic modification, as detailed below. IL-3 and hydrocortisone were removed on day 8 and SCF was removed on day 12. Cells were replated at ∼1×10^4^ cells/ml on day 5, <5×10^5^ cells/ml on day 8, and 7.5×10^5^-1.0×10^6^ cells/ml on day 12. The cells were co-cultured at 1×10^6^ cells/ml on a murine stromal cell layer (MS-5) beginning ∼day 14 ^38, 39^, and harvested between days 18-20. Growth and differentiation were monitored by hemocytometer and light microscopy of cytospin preparations stained with May-Grünwald and Giemsa. Enucleation rate was quantified using Vybrant DyeCycle violet (Life Technologies) (1:10,000) at 37° C for 30 min, followed by flow cytometry analysis on a MACSQuant flow cytometer (Miltenyi).

### Genetic modification of primary human CD34^+^ cells via CRISPR/Cas9

Two sgRNAs targeting human *CD44* were designed using the GPP sgRNA design portal (Broad Institute) and synthesized as chemically modified sgRNAs by Synthego. CD44-Cr1 (GAATACACCTGCAAAGCGGC) is predicted to recognize a sequence in exon 2 and CD44-Cr2 (GCAATATGTGTCATACTGGG) is predicted to recognize a sequence in exon 3. RNPs containing 300 pmol of each sgRNA and 150 pmol recombinant Cas9-NLS protein (Berkeley Macrolab) were introduced into 1×10^5^ CD34^+^ cells using a Lonza Amaxa 4-D nucleofector with program EO-100. Knock-out efficiency was determined after day 9 by flow cytometry.

### Enrichment for CD44-null erythroid precursors

Day 9-11 erythroid precursors were incubated with anti-CD44 antibody-immobilized beads (Miltenyi) (10µl beads for 6×10^6^ cells) at 4° C for 15 min. The cells were passed through an LD magnetic column (Miltenyi), the flow through was collected, and percent of CD44-null cells was determined by flow cytometry. A second round of enrichment was performed if CD44-null percent was less than 95%. After, cells were washed and cultured in PIMDM media as detailed above.

### Invasion assays

Parasite invasion assays were performed using late-stage schizont parasites isolated by magnet purification and added at 1.0-1.5% initial parasitemia to cultured red blood cells (cRBCs) or peripheral red blood cells (pRBCs) at 0.3% hematocrit in a volume of 100 μl per well in a 96 well plate. The ring-stage parasitemia was determined after 18-24 hours by bright-field microscopy of cytospin preparations stained with May-Grünwald and Giemsa. A minimum of 1000 cells were counted blindly for each technical replicate. For the CD44-CRISPR assays, invasion efficiency was determined for each of four biological replicates by normalizing the ring stage parasitema in each genetic background (mean of three technical replicates) to the average ring stage parasitemia in control pRBCs. For the invasion assays using CD44-null cRBCs enriched by magnet purification, invasion efficiency was determined for each of four biological replicates by normalizing the ring stage parasitemia in the CD44-null cells to the ring stage parasitemia in the isogenic control WT cRBCs (each biological replicate was an average of three technical replicates).

### Free merozoite binding assays

Free merozoites were isolated as previously described, with minor modifications ^40^. *P. falciparum* strain 3D7 schizonts were collected by magnet purification, incubated in 10 µM E64 for 7-8 hours, and released by syringe lysis. Hemozoin was removed by running lysate through an LS column (Miltenyi) on a MACS magnet. 12 µl (3 µg) of rCD44-His (SinoBiological 12211-H08H), rGYPC-His (SinoBiological 15627-H07H), rCD99-Fc (Creative Biomart CD99-473H), rCD44-Fc (SinoBiological 12211-H02H) or mock were incubated with 200µl of merozoite suspension for one hour at room temperature (RT), then incubated in 100µl anti-6X-his mouse monoclonal antibody (1:300), anti-CD99 (1:250) or anti-CD44 BRIC 222 (1:1000) for 1 hour, followed by goat anti-mouse IgG-FITC (1:500) for 1 hour. Binding of recombinant proteins to merozoites was measured by flow cytometry and data were analysed by FlowJo (v.10.6.1).

### Immunofluorescence Assays

Free merozoites were fixed in cold 4% paraformaldehyde/0.0075% glutaraldehyde in PBS for 10 minutes, resuspended in 1ml 0.3% BSA in PBS, and incubated with 2µg of rCD44-Fc (SinoBiological 12211-H02H) for one hour at RT. After washing, the merozoites were transferred to poly-L-lysine cover slips, blocked in PBS with 3% BSA, and incubated in primary antibodies anti-RAP1/2 mouse monoclonal (1:500) and anti-CD44 rabbit polyclonal (1:500) followed by goat anti-mouse Alexa Fluor 488 (1 µg/ml) and goat anti-rabbit Alexa Fluor 555 (4 µg/ml). Coverslips were mounted in Fluoromount-G with Dapi (Thermo). Images were acquired on a Keyence BZ-X fluorescence microscope.

### Pull-down assays

Schizont-stage lysate from ∼300ml *P. falciparum* strain 3D7 was prepared in ice cold PBS with protease inhibitors (PI; Pierce), addition of 1X saponin and incubation on ice for 5 minutes. Parasite pellets were washed several times in PBS with PI and resuspended in 500 µl lysis buffer (50 mM Tris/HCl Ph 7.5, 150 mM NaCl, 1% NP-40 in PBS with PI) on ice for 15-20 min with agitation, followed by centrifugation for 10 min at 4° C to recover supernatant. The NP-40 concentration in the parasite lysate supernatant was then diluted to 0.25% NP-40 by adding additional buffer (50 mM Tris/HCl Ph7.5 and 150 mM NaCl). For pull-down assays, 8 µg of rCD44-Fc (R&D Systems 3660-CD) or rIgG-Fc (R&D systems 110-HG) were incubated with 50 µl of protein A Dynabeads (Invitrogen) for one hour at room temperature. The beads were then resuspended in ∼500 µl parasite lysate and incubated overnight at 4° C. Bound proteins were eluted and separated by gel electrophoresis prior to submission to the Stanford University Mass Spectrometry facility (SUMS) for protein identification. In a typical mass spectrometry experiment, protein gel samples were destained, diced into 1 mm cubes, reduced in 5 mM DTT at 55C for thirty minutes, and then alkylated with 10 mM acrylamide for thirty minutes at RT to modify cysteines. The samples were then digested with trypsin/LysC (Promega) in the presence of ProteaseMax (Promega) overnight, before extraction and drying prior to LC/MS analysis. Mass spectra were collected on an Orbitrap Elite (RRID:SCR_018694 Thermo Scientific) coupled to a nanoAcquity Liquid Chromatograph (Waters Corporation). The mass spectrometer was operated in a data-dependent fashion using CID fragmentation in the ion trap for MS/MS spectra generation. The collected mass spectra were analyzed using Byonic v2.14.27 (Protein Metrics) as the search engine for peptide identification and protein inference. The precursor mass tolerances were set to 12 ppm, with 0.4 Da tolerances for MS/MS spectra. The search assumed semitryptic digestion and allowed for up to two missed cleavages against a database containing both *P. falciparum* and *Homo sapiens* sequences, as well as common contaminants. Data were validated using the standard reverse-decoy technique at a 1% false discovery rate using standard approaches. Data from the two bands submitted for CD44-Fc and for IgG-Fc were condensed using ComByne software to facilitate comparative analysis, generating a new coverage percentage and total number of spectra for each. To obtain a ranking of protein candidates enriched in rCD44-Fc over rFc, we used a cut off of >2-5 Logprob, >2 spectral counts, and calculate the NSAF (the number of spectral counts identifying a protein divided by the protein’s length divided by the sum of all spectral counts/lengths for all proteins in the experiment).

For immunoblotting, eluate fractions were run on an SDS-page gel and transferred for blotting with anti-EBA-175 rabbit anti-serum (1:2000), or anti-EBA-140 rabbit anti-serum (1:2000) in PBS-T with 5% BSA overnight at 4°C followed by goat anti-rabbit-HRP secondary antibody (1:25,000) for one hour, followed by detection with SuperSignal West Pico chemiluminescent substrate (Thermo).

### Recombinant protein expression and purification

RII-EBA-175-His and RII-EBA-140-His were expressed and purified as described previously ^12, 14, 15, 41^. Briefly, overnight cultures of *E. coli* Rosetta (DE3) cells were transfected with plasmid for either recombinant protein and grown at 37°C. 1L of luria-broth was then inoculated with the overnight culture and grown at 37°C until the cultures reached an O.D.600 of 0.6, at which time, expression was induced by adding 0.1mM isopropyl-1-thio-ý-galactopyranoside. Protein production was then allowed to proceed for 4 hours at 37°C. Cells were harvested by centrifugation and lysed and sonicated in 50mM Tris pH 8.0, 100mM NaCl, 5mM dithiothreitol, protease inhibitor tablets, and 1mg/mL lysozyme. Protein was extracted from inclusion bodies using 6M guanidine hydrochloride, 50mM Tris pH 8.0, 100mM NaCl, and 5mM dithiothreitol. The denatured protein was refolded by diluting dropwise into 400 mM l-arginine, 50 mM tris pH 8.0, 10 mM ethylenediaminetetraacetic acid, 0.1 mM phenylmethanesulfonyl fluoride, and either 2 mM reduced glutathione/2 mM oxidized glutathione for RII-EBA-175-His or 2 mM reduced glutathione/0.2 mM oxidized glutathione for RII-EBA-140-His and allowing the protein to refold for 48 hours at 4°C. The refolded proteins were then purified in batch by ion exchange resin followed by size exclusion chromatography using a HiLoad Superdex 200 16/60 column (Cytiva) and phosphate buffered saline (PBS).

### Recombinant protein binding assays

Protein A Dynabeads (1.5 mg) were incubated with 8 µg rCD44-Fc (R&D systems 3660-CD), rCD44-Fc (SinoBiological 12211-H02H), rCD55-Fc (SinoBiological 10101-H02H), IgG_1_-Fc (R&D systems 110-HG), or no protein in 200 µl antibody binding and washing buffer (Invitrogen kit 10006D) for one hour at RT. Recombinant proteins from SinoBiological were made in HEK293 cells and R&D proteins were made in a mouse myeloma cell line. The protein-coated beads were then incubated with 2 µM RII EBA-175-His or RII-EBA-140-His in PBS for one hour at RT. In antibody blocking experiments, anti-CD44 BRIC 222 (IBGRL) or control antibody BRIC 170 (targeting cytoplasmic domain of Band3) were incubated with the rCD44-Fc beads at 50 µg/ml for one hour prior to incubation with the EBA proteins. After washing, beads were incubated in primary antibody mouse anti-6x-His IgG (1:500) for 1 hour at RT and then secondary antibody goat anti-mouse FITC (1:1000). Binding was detected by flow cytometry (MACSQuant) and data were analyzed using FlowJo (v.10.6.1). For experiments to determine if binding was dependent on sialic acid, 10 µg rCD44-Fc was treated with 0.5 µl neuraminidase in 50 µl reaction volume for 1.5 hours at 37° C prior to binding to protein A beads. Glycoprotein staining was performed using the Pierce Glycoprotein Staining Kit (Thermo 24562), which included positive and negative controls HRP and STI, respectively.

### HUDEP-2 cell culture, differentiation, and genetic modification

The HUDEP-2 cell line was kindly provided by Drs. Ryo Kurita and Yukio Nakamura ^42^. Cells were maintained in Stem Span (StemCell Technologies) with 100 ng/ml SCF, 3 IU/ml Epo, 395 ng/ml dexamethasone (dex), and 1 µg/ml doxycycline (dox). The cells were split every 2-3 days to maintain concentration <1×10^6^ cells/ml. To induce erythroid differentiation, HUDEP-2 were washed and plated in PIMDM media with 5% Octaplas, 3 IU/ml Epo, and 1 µg/ml dox at 7×10^5^ cells/ml. After two days, dox was removed and the cells were plated at 7×10^5^ cells/ml. They were co-cultured on MS-5 stromal cell layer in PIMDM with Epo, 10% Octaplas and 0.5% F-68 at 1e6 cells/ml. Orthochromatic cRBCs were recovered on a Percoll gradient consisting of 2.5 ml 50% Percoll and 2.5 ml 55% Percoll.

To generate GYPA-null cells, HUDEP-2 were transduced at an MOI <1 with lentivirus prepared from pXPR_101 (lentiCas9-blast, gift from Feng Zhang, Addgene plasmid 52962) and selected with blasticidin. The cells were then transduced with lentivirus prepared from plentiguide-puro-GYPA-CR2 which was made by cloning annealed oligos GPA-C2F (CACCGTACCGGTTTCCTCTTCTGGA) and GPA-C2R (AAACTCCAGAAGAGGAAACCGGTAC) into pXPR003 (pLentiguide-puro, gift from Feng Zhang, Addgene plasmid 52963) at the *BsmB1* site. Cells were selected with puromycin and cloned by limiting dilution. Absence of GYPA surface expression was confirmed by flow cytometry. To generate the GYPC-null and CD44-null HUDEP-2 cells, RNP complexes consisting of rCas9 and two chemically modified sgRNAs targeting GYPC or one chemically modified sgRNA targeting CD44 (Synthego) were introduced into HUDEP-2 cells by nucleofection as above. GYPC-CR1 has the sequence CAUCCAGACAUCCCUGGAUC. GYPC-CR2 has the sequence AAUGUCCAUUAUGGUGGGGG. CD44-CR1 has the sequence CGUGGAAUACACCUGCAAAG. Clones were generated by limiting dilution and confirmed to be negative for GYPC or CD44 surface expression by flow cytometry. Double mutants (GYPA-null/CD44-null and GYPC-null/CD44-null) were generated by nucleofection of GYPA-null HUDEP-2 or GYPC-null HUDEP-2 cells with CD44-CR1, as above. Double-mutant clones were selected by FACS and confirmed by flow cytometry and sequencing.

### HUDEP-2 cRBC binding assays

1 ×10^6^ HUDEP-2 cRBCs differentiated to the orthochromatic erythroblast stage were washed in blocking buffer (DMEM with 10% FBS and 100 mM NaCl) and then incubated in recombinant RII EBA175-His or RII EBA-140-His at 5 µM final concentration. After washing, samples were incubated in anti-6x-His mouse monoclonal antibody (1:300) for one hour, and then goat anti-mouse IgG FITC (1:1000). Binding of recombinant proteins to the cRBCs was measured by flow cytometry and data were analyzed by FlowJo (v.10.6.1).

### Proteomic analysis and mass spectrometry

Isogenic WT and CD44-CRISPR cRBCs were generated from primary human HSPCs, and CD44-null cRBCs were isolated using microbeads, as above. Day 20 cRBCs were resuspended at 50% hematocrit in Krebs-Ringer buffer. For each simulation, 2×10^8^ cells at 50% hematocrit were incubated with PBS (mock) or RII EBA-175 (2 µM final concentration) for 30 minutes with agitation. Cells were then washed twice in cold Krebs-Ringer buffer, and resuspended in 1.4ml ice-cold lysis buffer (5 mM sodium phosphate pH8), Pierce protease inhibitors (Thermo A32963), and 0.5ml Halt phosphatase inhibitor (Thermo 78420). Samples were incubated on ice for 10 minutes to allow complete lysis, then centrifuged at 21,000 x G for > 25 min at 4°C. The ghost membrane layer was washed several times with 900 µl ice-cold lysis buffer, snap frozen, and sent to Applied Biomics, Inc. for 2D-DIGE analysis (Hayward, CA, USA). The samples were labelled with fluorescent dyes (Cy2, Cy3, or phospho-tag) and run on 2D-DIGE to separate proteins by size and pH, as per the vendor’s specifications (Gel 1: WT mock vs EBA-175 stimulated; Gel 2: CD44-null mock vs EBA-175 stimulated; Gel 3: WT EBA-175 stimulated vs phospho-stained; Gel 4: CD44-null EBA-175 stimulated vs phospho-stained; Gel 5: WT mock vs phospho-stained; Gel 6: WT EBA-140-stimulated vs phospho-stained; Gel 7: CD44-null mock vs phospho-stained; Gel 8: CD44-null EBA-140 stimulated vs phospho-stained). In-gel analysis generated gel images of individual samples and overlays of two samples. DeCyder software was used by Applied Biomics for quantitative analysis and comparison of all spots between the different samples, yielding quantitative protein expression ratios. Spot picking for mass spectrometry identification was performed by Applied Biomics based on differential intensity in stimulated versus unstimulated samples in WT vs CD44-null backgrounds. Protein identification was based on peptide fingerprint mass mapping (using MS data) and peptide fragmentation mapping (using MS/MS data). The MASCOT search engine was used to identify proteins from primary sequence databases.

### Data analysis

The DeCyder analysis was performed by Applied Biomics using the DeCyder 2D software from GE Healthcare, v.6.5: http://biotech.gsu.edu/core/Documents/Manuals/decyder-v6.5-manual.pdf. For statistical analysis of the invasion assays, two-tailed student t-tests were performed using GraphPad Prism v.9.

### Data Sharing Statement

For original data, please contact. Proteomics data from the 2D-DIGE and pull-down experiments may be found in data supplements available with the online version of this article. Raw proteomics data can be found at the PRIDE repository.

## RESULTS

### Efficient generation of CD44-null cRBCs from primary human HSPCs

To begin to characterize the role of CD44 in *P. falciparum* invasion, we sought to generate CD44-null RBCs using CRISPR/Cas9 genome editing in primary human hematopoietic stem/progenitor cells (HSPCs) followed by ex-vivo erythropoiesis (Fig. 1A). Nucleofection of ribonucleoprotein complexes containing two CD44-targeting sgRNAs and recombinant Cas9 resulted in an efficiency of CD44 knockout of ∼75%, as measured by flow cytometry (Fig. 1B). Both CD44-CRISPR and control WT populations of erythroid progenitors demonstrated a high rate of proliferation, with ∼10,000-fold increase in cell number over 15 days (Fig. 1C). After induction of terminal erythroid differentiation, the CD44-CRISPR cells developed in a similar manner to the control WT cells, progressing through morphologically distinct basophilic, polychromatic, and orthochromatic erythroblast stages (Fig. 1D). Importantly, they also underwent efficient enucleation, reproducibly generating >95% enucleated cRBCs (Fig. 1D-E). Together, these findings indicate that CD44 can be efficiently deleted from primary human HSPCs, and is not required for *ex-vivo* erythropoiesis, terminal erythroid differentiation, or enucleation.

**Figure 1.**
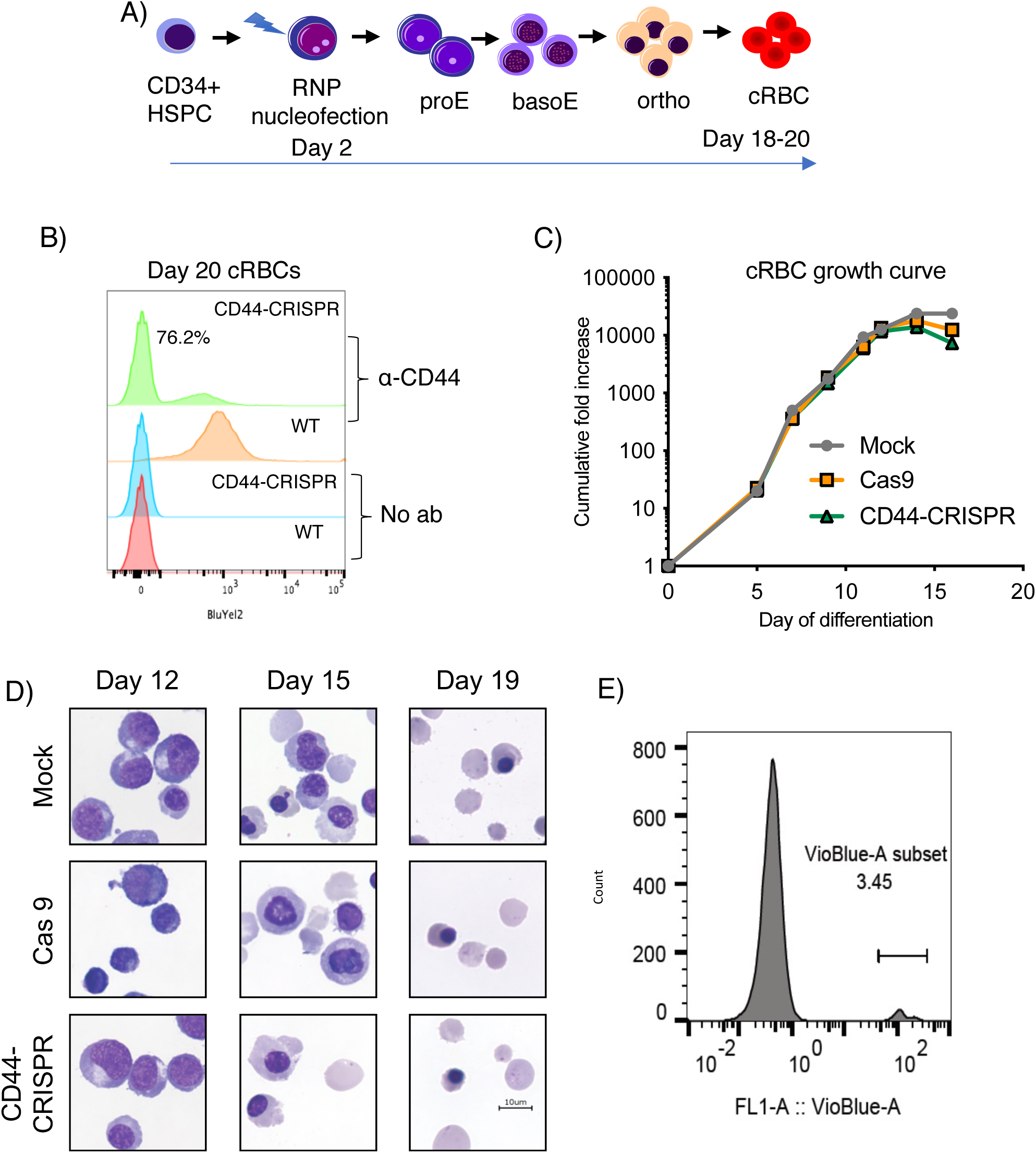
CD44 is dispensable for erythroid differentiation of primary human HSPCs. (A) Schematic of genome editing strategy to generate CD44-null cRBCs from primary human CD34^+^ HSPCs using CRISPR/Cas9. Ribonucleoprotein (RNP) complexes consisting of rCas9 and two CD44-targeting sgRNAs were introduced by nucleofection on day 2 of culture. Differentiating erythroblasts were plated on a murine stromal cell layer on day ∼13-14 to facilitate recovery of enucleated cRBCs. (B) Flow cytometry analysis of CD44 surface expression in unmodified (WT) cRBCs, or an isogenic population that was nucleofected with RNPs targeting CD44 (CD44-CRISPR). (C) Representative growth curves of primary human CD34^+^ HSPCs during ex-vivo erythropoiesis. Nucleofection was performed on day 2 to generate CD44-CRISPR cells, Cas9 control cells (WT cells nucleofected with rCas9 only), or Mock control cells (non-transfected). (D) Cytospin images of CD44-CRISPR versus isogenic WT erythroid cells during timecourse of ex-vivo erythropoiesis. Cas9, nucleofected with Cas9 only. Mock, non-transfected. (E) Representative experiment showing enucleation rate of CD44-null cRBCs upon terminal differentiation, as detected by flow cytometry using the cell-permeable DNA stain Vybrant DyeCycle Violet.

### Impaired *P. falciparum* invasion into CD44-null cRBCs

To investigate the requirement for erythrocyte CD44 in *P. falciparum* invasion, we performed invasion assays using isogenic WT and CD44-CRISPR mutant cRBCs generated from primary human CD34^+^ HSPCs by CRISPR/Cas9 and ex-vivo erythropoiesis, as described above. *P. falciparum* strain 3D7 was synchronized to the schizont stage and incubated with the CD44-CRISPR or isogenic WT cRBCs for ∼18 hours to allow for egress and invasion, and the resulting parasitemia was quantified by blinded counts of cytospin slides. The results showed that the parasitemia in the CD44-CRISPR population was reduced by ∼50% relative to the WT control cRBCs, confirming that *P. falciparum* infection is impaired in the absence of CD44 (p=0.0062; Fig. 2A). Visual inspection of the ring-stage parasites growing in WT versus CD44-CRISPR cRBCs did not reveal any gross morphological differences between the two genetic backgrounds (Fig. 2B). While most infected cells harbored one or two parasites, occasional cells with multiple rings could be detected in both CD44-CRISPR and isogenic WT cRBCs.

**Figure 2.**
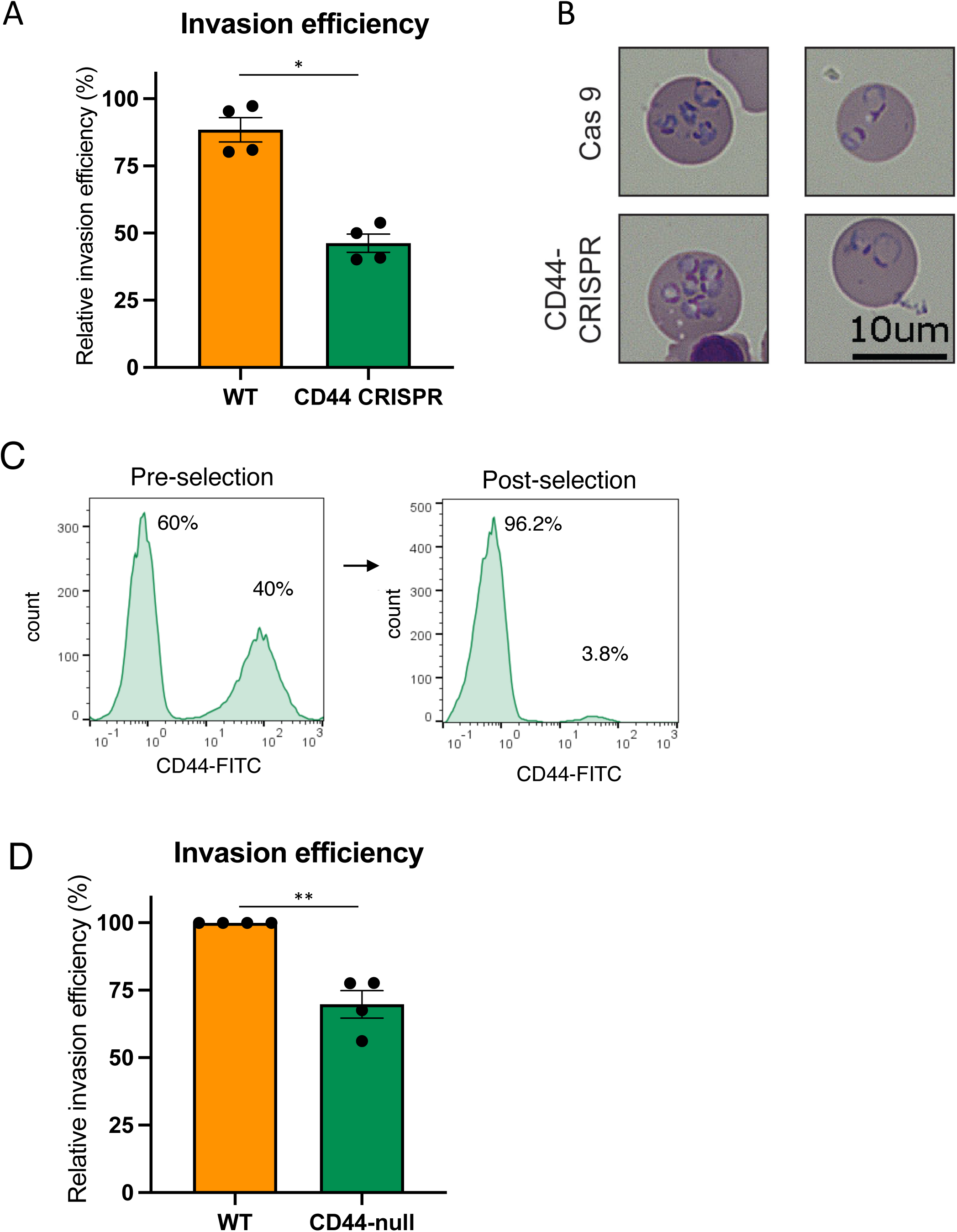
CD44 is required for efficient *P. falciparum* invasion. (A) Invasion assays of *P. falciparum* strain 3D7 into WT cRBCs versus CD44-CRISPR cRBCs. Approximately 18 hours after incubation with late-stage schizonts, parasitemia was determined by blinded counting of cytospin slides stained with May-Grünwald and Giemsa, and is presented relative to parasitemia in control erythrocytes. Plotted are four biological replicates, each of which was performed in triplicate, with the line indicating the mean ± S.E.M. *, p = 0.0062, two-tailed paired t-test. (B) Cytospin images of *P. falciparum* parasites ∼18 hours after invasion into isogenic WT or CD44-CRISPR cRBCs. (C) Flow cytometry plot showing efficiency of negative selection for CD44-null cRBCs using CD44 microbeads. (D) Invasion assays of *P. falciparum* strain 3D7 into WT cRBCs versus CD44-null cRBCs isolated by negative selection. Parasitemia was determined by blinded counting of cytospin slides by two individuals, and is presented normalized to the parasitemia in the WT cRBCs for each replicate. Each of four biological replicates are plotted, each of which were performed in triplicate, with the line indicating the mean of the biological replicates, ± S.E.M. **, p=0.0010, two-tailed t-test.

The invasion phenotype observed in the CD44-CRISPR population was reminiscent of prior results showing that *P. falciparum* invasion was reduced by ∼50% in CD44-knockdown (KD) cRBCs derived from primary human HSPCs and in CD44-null jkRBC erythroblasts derived from the erythroleukemic line JK-1 ^20, 21^. To determine if CD44 is essential for *P. falciparum* invasion, we used negative selection to isolate a purer population of CD44-null cRBCs for use in invasion assays (Fig. 2C). Again, we observed a moderate invasion phenotype in the CD44-null cRBCs relative to isogenic control cells (p=0.0010; Fig. 2D). Together, these results confirm that CD44 is necessary for efficient *P. falciparum* invasion of erythrocytes, but suggest that it is not essential for invasion.

### Direct interaction between *P. falciparum* merozoites and erythrocyte CD44

Given that CD44 has a large extracellular domain, is an established pathogen receptor on epithelial cells, and is required for efficient *P. falciparum* infection of cRBCs, we hypothesized that it may act as a receptor for parasite invasion. To investigate this, we isolated free merozoites and quantified their interaction with recombinant CD44 using flow cytometry-based assays (Fig. 3A-B). Recombinant GYPC and recombinant CD99 served as positive and negative controls, respectively, as GYPC is a known receptor for *P. falciparum*, and available evidence suggests CD99 has no role in *P. falciparum* invasion ^8, 20^. Free merozoites were observed to bind directly to rCD44-Fc but not to rCD99-Fc, confirming that the interaction observed with rCD44-Fc was not attributable to the Fc domain (Fig. 3A). Binding was also observed for rCD44-His and the positive control rGYPC-His, as measured using an anti-His antibody (Fig. 3B). To further investigate the interaction between CD44 and merozoites, we performed immunofluorescence assays to determine the localization of rCD44 bound to free merozoites. CD44 localized to the apical end of merozoites, as indicated by its co-localization with the known apical marker RAP1 (Fig. 3C). Notably, many *P. falciparum* invasion ligands are known to be concentrated at the apical end of the merozoite, from which invasion commences. Together, these results suggest that CD44 can act as a receptor for invasion by interacting with a parasite ligand localized in the apical region of the merozoite.

**Figure 3.**
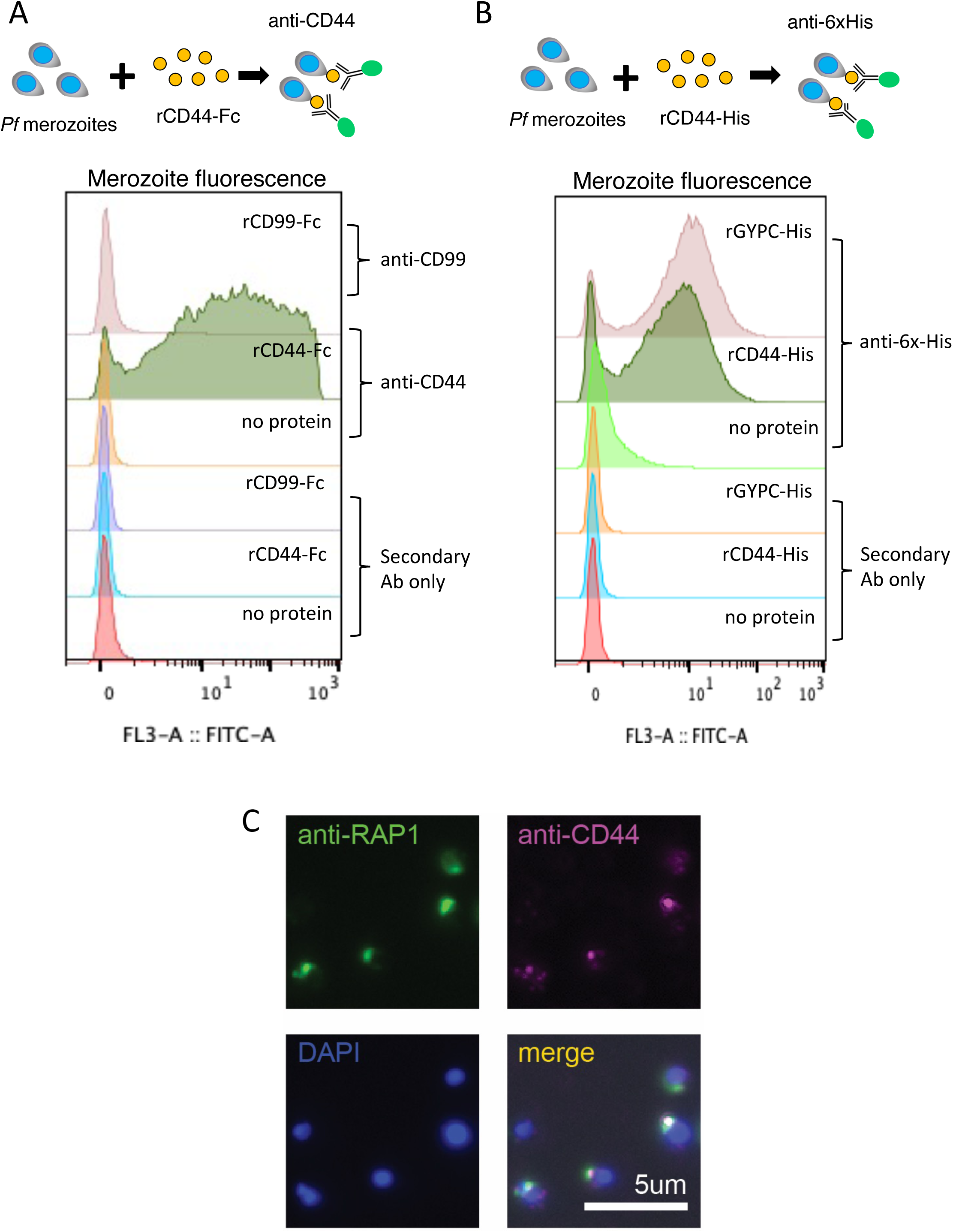
*P. falciparum* merozoites bind to CD44. (A) Flow cytometry-based assay to quantify binding between free *P. falciparum* merozoites and recombinant CD44-Fc or CD99-Fc proteins. Free merozoites were incubated with recombinant proteins, and binding was detected using anti-CD44 or anti-CD99 antibodies. (B) Flow cytometry-based assay to quantify binding between free *P. falciparum* merozoites and recombinant CD44-His or GYPC-His proteins. Histograms depicting merozoite fluorescence upon incubation with recombinant His-tagged proteins and detection with anti-His antibody. (C) Images of immunofluorescence assays of free *P. falciparum* merozoites after binding to rCD44-His, and its co-localization with the merozoite protein RAP1. Images were taken with a Keyence BZ-X700 all-in-one fluorescence microscope at 100x magnification under oil immersion.

### CD44 interacts with the *P. falciparum* invasion ligands EBA-175 and EBA-140

To identify *P. falciparum* ligand(s) that interact with CD44, we performed *in vitro* pull-down assays by incubating rCD44-Fc beads or rIgG_1_-Fc (rFc) beads with lysate from *P. falciparum* schizont-stage parasites, followed by mass spectrometry. Analysis of the mass spectrometry data from the eluates revealed a strong enrichment for two *P. falciparum* invasion ligands from the Erythrocyte Binding Antigen (EBA) family in the rCD44-Fc pull-down relative to the rFc control condition: EBA-175 and EBA-140 (Fig. 4A and Supplementary Dataset 1). In addition, the large number of unique peptides detected and the percent coverage of these proteins indicated an efficient pull-down (Fig. 4A). These results were confirmed by independent pull-down experiments followed by western blots, showing that EBA-175 and EBA-140 were specifically detected in the elution from the pull-down using rCD44-Fc but not rFc (Fig. 4B). These findings demonstrate that CD44 can interact with EBA-175 and EBA-140, either directly or indirectly.

**Figure 4.**
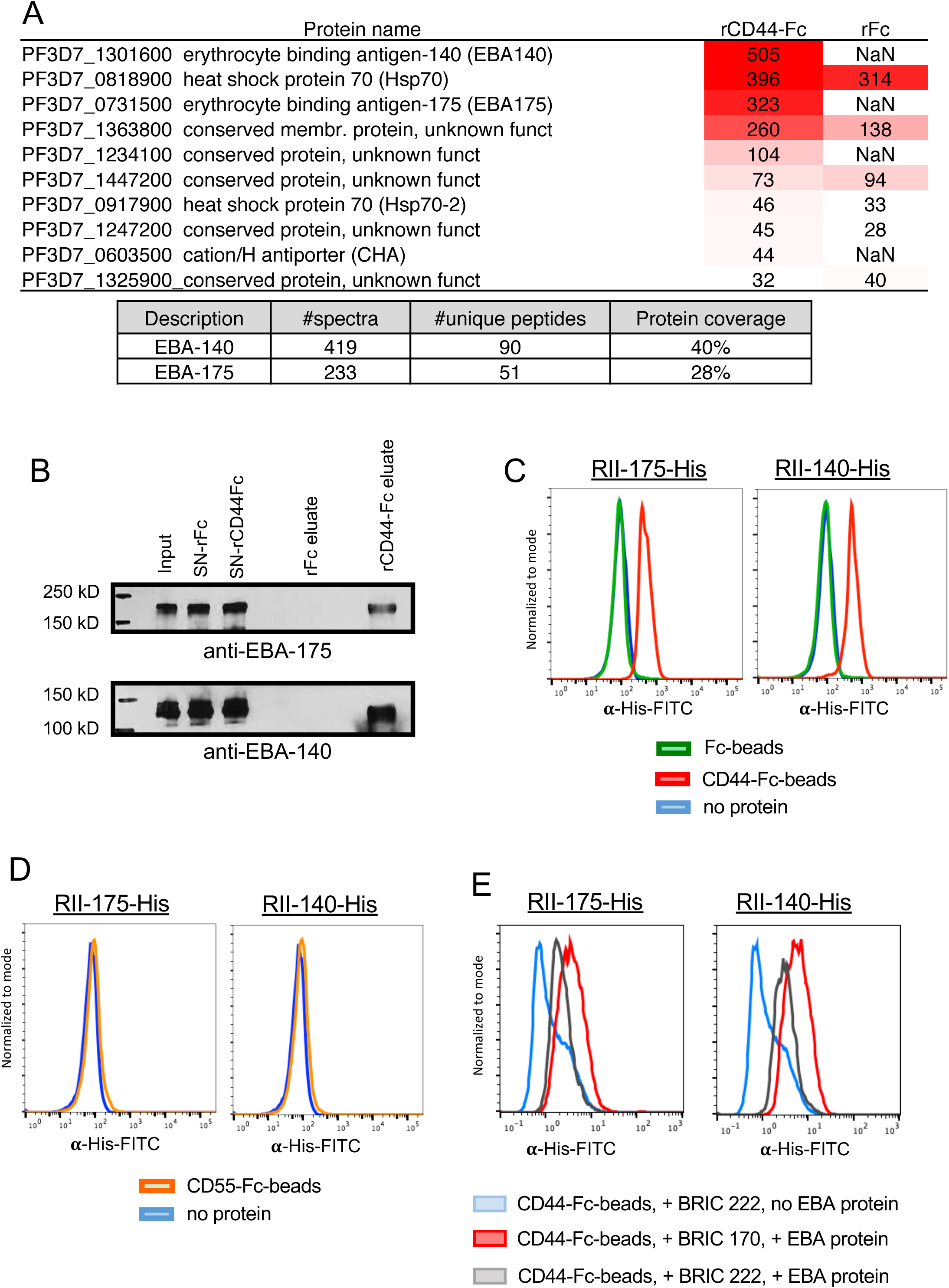
CD44 specifically interacts with *P. falciparum* invasion ligands EBA-175 and EBA-140. (A) Mass spectrometry results from affinity purification experiments in which bead-immobilized rCD44-Fc or rFc were incubated with lysate from *P. falciparum* strain 3D7 schizont-stage parasites. Heat map shows the normalized spectral counts (based on protein size) of the most abundant *P. falciparum* proteins detected and their enrichment in rCD44-Fc compared to rFc. NaN indicates no counts. The table shows Log probability, raw number of spectra and unique peptides, and coverage of EBA-140 and EBA-175 proteins found in rCD44-Fc lane. (B) Western blot of independent affinity purification experiment in which bead-immobilized rCD44-Fc or rFc were incubated with *P. falciparum* schizont-stage lysate, followed by immunoblotting for EBA-175 or EBA-140. SN, supernatant. (C) Flow cytometry-based binding assays in which recombinant region RII of EBA-175-His (left) or EBA-140-His (right) were incubated with beads coated with rCD44-Fc, rFc, or no protein. Binding was detected using an anti-His antibody and a fluorescent secondary antibody. (D) Flow cytometry-based binding assays between rCD55-Fc or rFc and region RII of EBA-175-His (left) or EBA-140-His (right). (E) Binding assays of RII EBA-175 (left) or RII EBA-140 (right) with rCD44-Fc in the presence of anti-CD44 monoclonal antibody BRIC 222 or isotype control BRIC 170.

Region II (RII) of EBA-175 and EBA-140 comprise the receptor binding domains ^2^. To determine if EBA-175 and EBA-140 RII interact with CD44 directly, we used rCD44-Fc and rFc-coated beads in *in vitro* binding assays with recombinant RII EBA-175-His and RII EBA-140-His. Binding of the RII-EBA proteins to the beads was measured by flow cytometry using an anti-His antibody. We observed a clear shift in fluorescence for rCD44-coated beads incubated with RII-175-His or RII-140-His, whereas no shift was seen for the rFc-coated beads or beads alone after incubation with the recombinant proteins, demonstrating direct interactions between rCD44-Fc and both EBA-140 and EBA-175 (Fig. 4C). Importantly, there was no detectable binding to CD55-Fc, another glycosylated erythrocyte membrane protein, further demonstrating the specificity of the observed interactions with rCD44-Fc (Fig. 4D and Fig. S1). Additionally, we observed that pre-incubation of the rCD44-Fc beads with anti-CD44 antibody BRIC 222 inhibited the binding by both EBA-175 and EBA-140, but this was not seen for isotype control antibody BRIC 170, which binds to an epitope in the cytoplasmic domain of Band3 (Fig. 4E). Together, these findings indicate that both RII-175 and RII-140 can interact directly and specifically with human CD44.

To determine if the binding of the EBA proteins to CD44 was dependent on sialic acid, we treated rCD44-Fc with neuraminidase (NM) to partially remove sialic acid. In *in vitro* binding assays, we observed a partial reduction in binding of EBA-175 and EBA-140 RII to NM-treated rCD44-Fc compared to untreated rCD44-Fc (Fig. S2). These results suggest that the interaction between the EBA proteins and rCD44 depends, at least in part, on sialic acid.

### CD44 is not a primary determinant of EBA-175 and EBA-140 binding to the RBC

Since the binding of EBA-175 and EBA-140 to RBCs is known to involve GYPA and GYPC, respectively, we next sought to determine how the interactions of EBA-175 and EBA-140 with CD44 may contribute to their RBC binding activity. To test this, we utilized the immortalized erythroid cell line HUDEP-2 ^42^, which is readily amenable to genetic manipulation and cloning, and which we found can be differentiated to orthochromatic erythroblasts permissive to *P. falciparum* invasion (Fig. 5A and Fig. S3).

**Figure 5.**
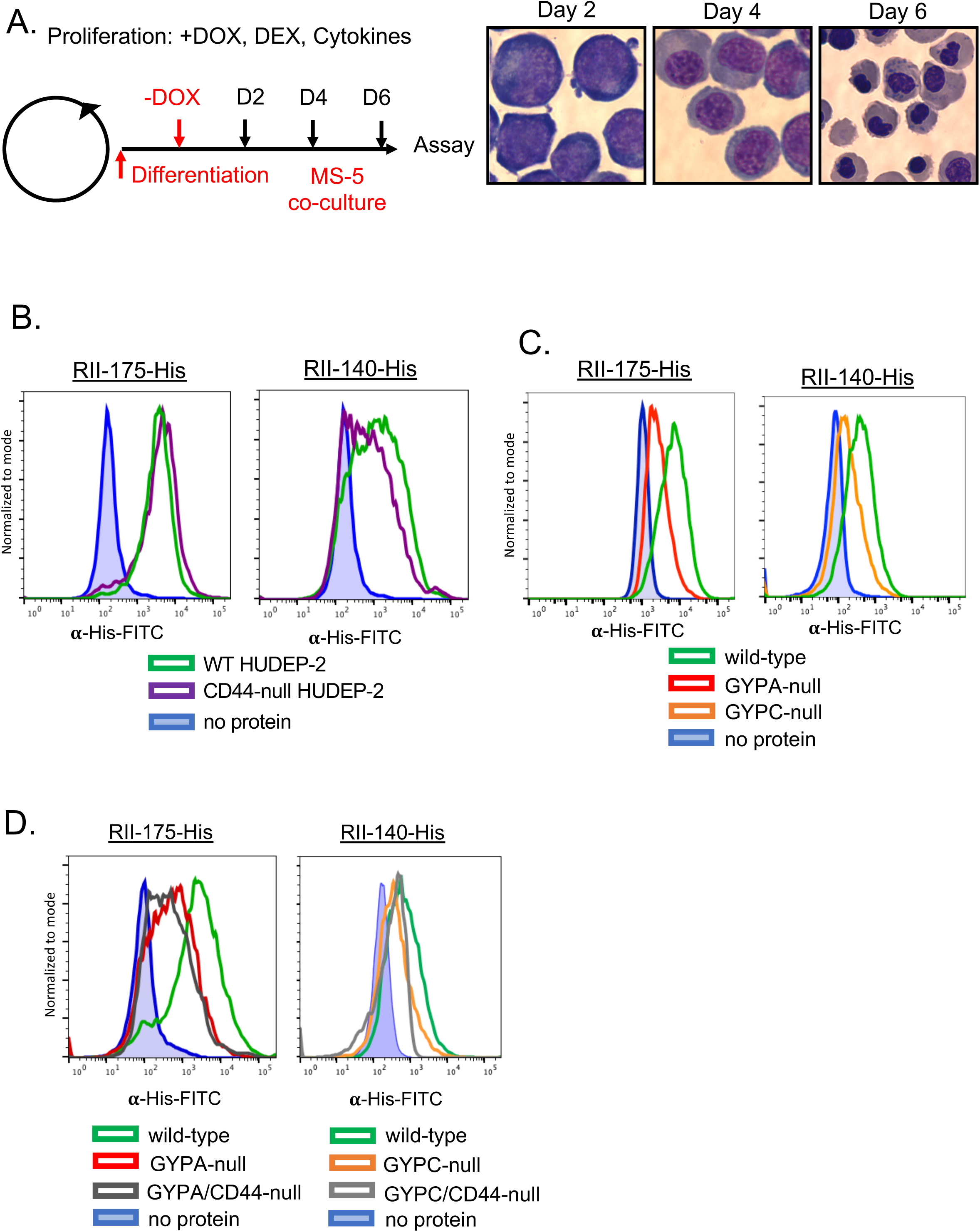
Genetic analysis in HUDEP-2 cells reveals that GYPA and GYPC are the primary determinants of EBA-175 and EBA-140 binding to cRBCs, respectively. (A) Schematic of HUDEP-2 proliferation and differentiation protocol, and representative images of differentiating HUDEP-2 cells. DOX, doxycycline; DEX, dexamethasone. Day 2, pro-erythroblasts; Day 4, polychromatic erythroblasts; Day 6, orthochromatic erythroblasts. (B) Flow cytometry-based binding assays of RII regions of EBA-175-His (left) and EBA-140-His (right) incubated with WT versus CD44-null HUDEP-2 orthochromatic erythroblasts. Binding of the EBA proteins to the cells was quantified using anti-His antibody and fluorescent secondary antibody. (C) Flow cytometry-based binding assays of RII regions of EBA-175-His (left) and EBA-140-His (right) incubated with WT, GYPA-null, or GYPC-null HUDEP-2 orthochromatic erythroblasts. Binding was quantified using anti-His antibody and fluorescent secondary antibody. (D) Flow cytometry-based binding assays of RII regions of EBA-175-His (left) and EBA-140-His (right) incubated with WT, GYPA-null, GYPA/CD44-null, GYPC-null, or GYPC/CD44-null HUDEP-2 orthochromatic erythroblasts. Binding was quantified using anti-His antibody and fluorescent secondary antibody.

We generated CD44-, GYPA-, or GYPC-null mutants in HUDEP-2 cells using CRISPR/Cas9 genome editing and confirmed that the mutant clones lacked surface expression of the respective proteins by flow cytometry (Fig. S4). After inducing differentiation down the erythroid lineage to orthochromatic erythroblasts (orthos), we used the cells in flow cytometry-based binding assays with recombinant RII-175-His and RII-140-His. The results showed that RII-175 bound to the WT and CD44-null orthos to a similar degree (Fig. 5B). Similar results were observed for RII-140, with only a slight reduction in binding noted for the CD44-null orthos (Fig. 5B). These findings suggest that CD44 is not a primary determinant of EBA-175 or EBA-140 binding to the HUDEP-2 orthos, particularly in comparison to the glycophorins. Indeed, in parallel experiments using GYPA-null or GYPC-null HUDEP-2 orthos, we observed that RII-175 and RII-140 binding was substantially dependent on GYPA and GYPC, respectively, consistent with these surface proteins being RBC receptors for EBA-175 and EBA-140 (Fig. 5C). Notably, CD44 is a surface protein of low abundance on human erythrocytes, with an estimated copy number of ∼10,000 molecules per cell, as compared to ∼1e6 copies of GYPA and ∼143,000 of GYPC ^4, 43^. Conceivably, this low copy number could impact the ability of CD44 to serve as prominent binding receptor relative to the glycophorins.

To determine whether the observed residual binding of the EBA proteins to GYPA-null and GYPC-null HUDEP-2 orthos was due to CD44, we next generated double mutant HUDEP-2 cells (GYPA/CD44-null or GYPC/CD44-null) and used them in binding assays. For RII-175, binding to the GYPA/CD44-null HUDEP-2 orthos was minimally reduced relative to GYPA-null HUDEP-2 orthos. Similarly, binding of RII-140 to GYPC/CD44-null HUDEP-2 orthos was slightly reduced relative to GYPC orthos (Fig. 5D). While these minor shifts are consistent with EBA-175 and EBA-140 being able to bind to CD44, they also confirm that CD44 is not a major determinant of the RBC binding activity of EBA-175 or EBA-140, relative to the glycophorins. Instead, these findings raise the hypothesis that CD44 may be acting as a co-receptor during *P. falciparum* invasion.

### A role for erythrocyte CD44 in EBA-175-induced signaling

In other cell types, CD44 has been shown to facilitate downstream signaling despite lacking intrinsic kinase activity ^28^. For example, CD44 acts as a co-receptor for the ErbB family of receptor tyrosine kinases and for the c-MET receptor, regulating diverse cellular processes ^35, 44–46^. As we found that CD44 can interact with EBA-175 and recent work has shown that EBA-175 binding to RBCs induces changed in membrane deformability and cytoskeletal protein phosphorylation ^47, 48^, we next investigated if CD44 plays a role in signaling in RBCs.

To determine if CD44 plays a role in EBA-175-induced signaling in the RBC, we generated pure populations of WT and isogenic CD44-null cRBCs for use in two-dimensional difference gel electrophoresis (2D-DIGE) experiments. Since 2D-DIGE can be performed with small amounts of material, this approach was more feasible than phosphoproteomics given the limitations of generating genetically-modified cRBCs from primary human HSPCs. We incubated WT or CD44-null cRBCs with recombinant EBA-175 or mock stimulus and performed 2D-DIGE on membrane ghost lysates to measure changes in protein intensity in the two genetic backgrounds, which would likely reflect post-translational modifications or protein stability since RBCs cannot perform protein synthesis (Fig. 6A). We identified 50 spots with ≥1.3-fold change in intensity in one or both genetic backgrounds after stimulation with EBA-175, as quantified by DeCyder analysis (Fig. 6B and Supplemental Dataset 2). Several proteins appeared to be differentially regulated by EBA-175 in WT vs CD44-null cRBCs, suggesting that CD44 may play a role in the regulation (Fig. 6C).

**Figure 6.**
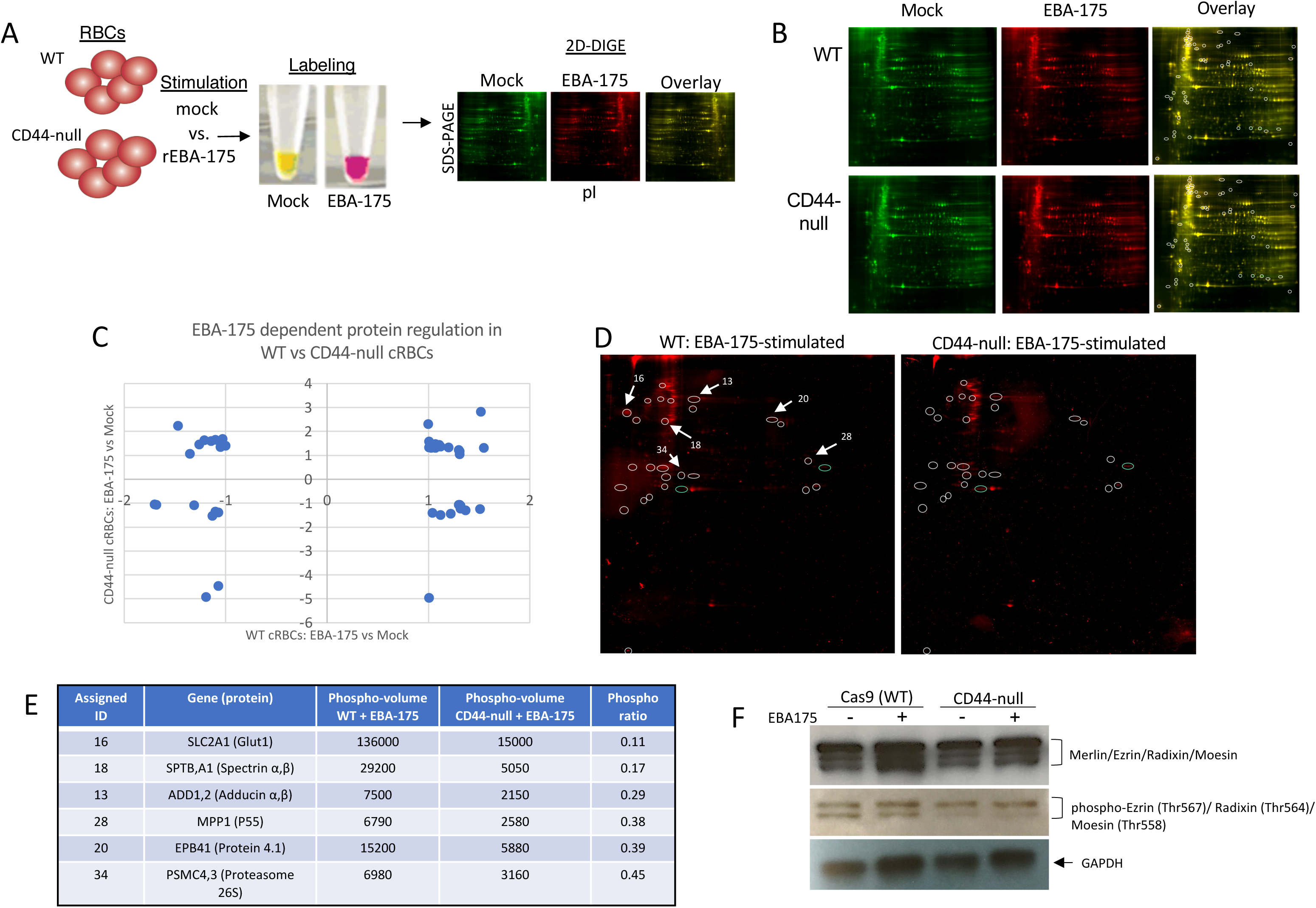
CD44 facilitates EBA-175-induced signalling to the RBC cytoskeleton. (A) Schematic of 2D-DIGE assays. (B) 2D-DIGE analysis of WT or CD44-null cRBC lysates stimulated with mock or RII EBA-175. Circles represent spots with ≥1.3-fold change in intensity in mock versus RII EBA-175-stimulated samples in one or both genetic backgrounds as identified by DeCyder analysis. (C) Quantification of the fold change in intensity in RII EBA-175-stimulated versus mock-stimulated WT cRBCs and RII EBA-175-stimulated versus mock-stimulated CD44-null cRBCs. Spots identified with >1.3-fold change in intensity were plotted. (D) 2D-DIGE analysis of protein phosphorylation in WT versus CD44-null cRBCs stimulated with RII EBA-175. The circles indicate 25 spots with ≥2-fold increase in phosphorylation in the stimulated WT cRBCs compared to CD44-null cRBCs. The six spots that were sent for identification by mass spectrometry analysis are indicated. (E) High-confidence proteins identified by mass spectrometry of 2D-DIGE spots with differential phosphorylation between EBA-175-stimulated WT versus CD44-null cRBCs. Phosphovolumes were quantified by Applied Biomics using DeCyder software. (F) Western blot for total and phosphorylated ERM complex proteins (merlin, ezrin, radixin, moesin) WT or CD44-null cRBCs stimulated with mock versus RII EBA-175. Anti-GAPDH was used as a loading control.

To determine if these shifts reflected altered protein phosphorylation, we performed 2D-DIGE on ghost lysates from EBA-175-stimulated WT or CD44-null cRBCs using a dye specific for phosphorylated proteins (Applied Biomics, Inc). This analysis identified 26 spots with ≥2-fold increased phosphorylation in the stimulated WT cRBC ghost lysate relative to the CD44-null background, indicating that EBA-175-induced phosphorylation of erythrocyte proteins is dependent on CD44, at least in part (Fig. 6D). We chose six of the most highly differentially phosphorylated spots between WT and CD44-null ghost lysates for protein identification by mass spectrometry. Interestingly, this analysis identified the cytoskeletal linker protein 4.1, as well as several components of the erythrocyte cytoskeletal network (SLC12A1, Adducin a/B, p55, and spectrin A/B) (Fig. 6E). These results are particularly intriguing because protein 4.1R is a cytoskeletal linker protein that regulates cell shape through its local protein-protein interactions, which include p55 ^49–51^, and possibly CD44 ^52, 53^. The ERM family members, ezrin, radixin, and moesin have also been shown to interact with CD44 as cytoskeletal linker proteins ^30, 31, 54^. While these proteins were not identified in our limited-scope mass spectrometry analysis, immunoblotting for the ERM proteins demonstrated reduced phosphorylation in CD44-null cRBCs relative to isogenic WT, consistent with a role for CD44 in signaling to the cytoskeleton (Fig. 6F).

Since EBA-140 can functionally substitute for EBA-175 to mediate invasion in certain strain backgrounds and we found that both proteins can interact with CD44, we also investigated RII EBA-140-induced RBC ghost lysate phosphorylation changes and their dependence on CD44 by 2D-DIGE. This analysis showed that stimulation with EBA-140 also leads to altered phosphorylation of several RBC proteins in a CD44-dependent manner, including ankyrin-1, spectrin alpha/beta, and protein 4.1 (Fig. S5).

## DISCUSSION

CD44 is recognized as a widely-expressed cellular adhesion molecule, impacting diverse cell processes such as migration, inflammatory responses, and metastasis, depending on the isoform and context. Its effects are exerted through interactions with ligands (e.g. hyaluronic acid), interactions with other surface receptors (e.g. c-Met), and interactions with cytoskeletal linker proteins through its conserved cytoplasmic domain ^28, 55^. Although CD44 has been extensively studied in leukocytes, epithelial cells and cancer cells, little is known about its function on human erythrocytes aside from its role in presenting the Indian blood group antigens ^22^. Recent evidence suggests that there may be an association between activation of the Gardos channel and the interaction of CD44 with HA, suggesting CD44 may impact erythrocyte senescence _56,57._

CD44 was first identified as a candidate host factor for *P. falciparum* though a pooled, forward genetic shRNA screen in cultured erythroblasts derived from primary human HSPCs ^20^. In the screen, CD44 was one of the top candidates identified, along with CD55, and its importance to *P. falciparum* was validated by the demonstration that invasion was reduced into CD44-knockdown enucleated cRBCs relative to isogenic WT cRBCs. Additional evidence for a role for CD44 in *P. falciparum* invasion was reported by Kanjee *et al*., where the rate of parasite invasion was reduced by ∼50% in CD44-knockout jkRBCs, as compared to isogenic WT jkRBCs at the same stage of development ^21^. In this work, we report the efficient generation of CD44-null cRBCs from primary human HSPCs using CRISPR/Cas9 genome editing for the first time. Importantly, the cRBCs were observed to proliferate and differentiate in a manner indistinguishable from isogenic WT cRBCs, demonstrating that CD44 is not required for erythroid development in human cells. Further, our finding that *P. falciparum* invasion was reduced but not completely inhibited in CD44-null cRBCs confirms that CD44 plays a role in *P. falciparum* invasion, but suggests it is not an essential host factor under these *in vitro* conditions.

Prior work in epithelial cells created precedent for the idea that CD44 can act as a pathogen receptor. Group A *Streptococcus* binds to CD44 on keratinocytes through its hyaluronic capsule, an interaction that triggers downstream RAC1-dependent signaling and cytoskeletal rearrangements in the host cell, enabling tissue invasion ^29^. CD44 on epithelial cells has also been shown to interact with the Shigella secreted protein IpaB, leading to cytoskeletal reorganization and increased invasion ^58^. CD44 has been best-studied as an adhesion molecule, where its interactions with hyaluronic acid or other components of the extracellular matrix can trigger a variety of downstream effects impacting cell growth, survival, and differentiation, depending on the context ^28^.

Reasoning that *P. falciparum* may exploit CD44 as a receptor for RBC invasion, we used pull-down assays to identify *P. falciparum* proteins that interact with CD44. The discovery of EBA-175 and EBA-140 as interactors of CD44 was surprising, as these well-established invasion ligands already have known receptors (GYPA and GYPC, respectively). Our observation that rCD44 can interact with free *P. falciparum* merozoites at their apical end supports this finding, as the EBA proteins are localized at the merozoite’s apical end. We confirmed the interactions using multiple methods, including immunoblotting, recombinant protein binding assays, and cellular binding assays using genetically modified erythroid cells. Of note, the cellular assays demonstrated that GYPA and GYPC are the primary determinants of RII EBA-175 and RII EBA-140 binding to RBCs, raising the hypothesis that CD44 may instead be acting as a co-receptor to facilitate invasion. Indeed, its low surface abundance on RBCs relative to the glycophorins and the relatively modest invasion phenotype (e.g. relative to basigin, which appears to be essential for *P. falciparum* invasion) could be consistent with a co-receptor function. Importantly, CD44 has been shown to function as a co-receptor with a variety of proteins, such as c-Met, EGFR, and TGF-β ^55^.

The observation that CD44 can interact with both EBA-175 and EBA-140 suggests it may be interacting with these proteins through a common domain. It is also consistent with prior work demonstrating that diverse *P. falciparum* strains rely on CD44 for efficient invasion, despite expressing different dominant invasion ligands ^21^. Future efforts aimed at mapping the critical resides for binding between CD44 and the EBA proteins will be important for detailed understanding of these interactions, and their relationship to the glycophorins. Interestingly, patient RBC samples deficient in protein 4.1R have been shown to have reduced surface expression of CD44 and GYPC, suggesting that these proteins may exist within a complex ^53^.

Our results provide evidence for a mechanism in which CD44 acts as a co-receptor for *P. falciparum* invasion. A co-receptor role could be consistent with the moderate invasion phenotype observed in the absence of CD44, at least compared to basigin, which is considered an essential receptor for parasite invasion ^17^. Co-receptors are generally defined as cell surface molecules that influence receptor-ligand activity but do not contain intrinsic catalytic activity. In the case of *P. falciparum* invasion, our results support a model where EBA protein binding to the RBC surface via the glycophorins also engages CD44, influencing the phosphorylation of cytoskeletal proteins via interactions between the cytoplasmic domain of CD44 and cytoskeletal linker proteins. Previous work has shown that EBA-175 binding to GYPA increases RBC deformability and is associated with changes in phosphorylation of cytoskeletal proteins ^47, 48^. EBA-175 shed from merozoites also induces RBC clustering and promotes parasite growth in culture ^41^. Since CD44 in an integral membrane protein that can coordinate adhesive and signaling events ^59^, our findings provide important insights into how *P. falciparum* EBA proteins can signal to the RBC cytoskeleton. Future work will be required to determine if CD44 is involved in the EBA-175-induced RBC clustering phenotype. A comprehensive understanding of the signaling pathways triggered during *P. falciparum* invasion will also require more extensive investigation, ideally via global phosphoproteomics and using genetically altered cRBCs stimulated with invading merozoites rather than individual invasion ligands.

As our understanding of host determinants of infectious diseases advances, novel, host-directed therapies that can counteract the evolution of pathogen resistance are beginning to be brought into clinical practice. This is an emerging concept in the malaria field, but increasing evidence suggests that targeting host factors can be an effective anti-malarial strategy ^60–62^. Given its role in several disease contexts, CD44 is currently being explored as a therapeutic target, particularly in the oncology field ^63, 64^. The demonstration in this work that *P. falciparum* exploits CD44 to facilitate RBC invasion lays a key foundation for future investigation into the potential of CD44-related pathways as targets for malaria intervention.

## ACKNOWLEDGEMENTS

We thank Nana Ansuah Peterson from the Egan Lab, Ryan Leib and colleagues from the Stanford University Mass Spectrometry Facility, and John Liao from Applied Biomics, Inc. (Hayward, CA) for technical assistance. We are grateful to members of the Egan Lab, John Boothroyd, Ellen Yeh, Paul Bollyky, and members of their labs for helpful discussions. We thank Ryo Kurita and Yukio Nakamura for contributing the HUDEP-2 cell line. Antibodies used in this work were kindly provided by Alan Cowman and Anthony Holder. Some mass spectrometry data were collected at the Vincent Coates Foundation Mass Spectrometry Laboratory, Stanford University Mass Spectrometry (RRID:SCR_017801). This work utilized the Thermo LTQ-Orbitrap Elite mass spectrometer system (RRID:SCR_018694) that was purchased with funding from National Institutes of Health Shared Instrumentation grant S10RR027425. This work was supported in part by NIH P30 CA124435 utilizing the Stanford Cancer Institute Proteomics/Mass Spectrometry Shared Resource. This work was supported in part by NIH DP2HL13718601 (E.S.E.) and a faculty scholar award from the Stanford Maternal Child Health Research Institute (E.S.E.). B.B., A.K.K., and M.T. were funded through postdoctoral fellowships from the Stanford Maternal Child Health Research Institute and A.K.K. was also supported by a T32 training grant in pediatric nonmalignant hematology and stem cell biology (T32DK098132-06A1). E.S.E. is a Chan Zuckerberg Biohub-San Francisco investigator and a Tashia and John Morgridge Endowed Faculty Scholar of the Stanford Maternal and Child Health Research Institute. N.H.T. and N.D.S are supported by the Intramural Research Program of the National Institute of Allergy and Infectious Diseases, National Institutes of Health.

## Author contributions

Conceptualization, B.B., C.Y.K., and E.S.E.; Performed experiments, B.B., C.Y.K., C.L., A.K., and E.S.E.; Data analysis, B.B., C.K., A.K., M.T., and E.S.E., Visualization, B.B., C.K., A.K., M.T., and E.S.E., Resources, N.H.T. and E.S.E.; Writing-original draft B.B. and E.S.E.; Writing-review and editing-B.B., C.Y.K., A.K.K., M.T., N.D.S., N.H.T., and E.S.E.; Supervision, E.S.E., Funding acquisition, E.S.E.

## Notes

### Competing Interest Statement

The authors have declared no competing interest.

