## Supplemental data for "*Plasmodium falciparum* exploits CD44 as a co-receptor for erythrocyte invasion"

### Figure S1

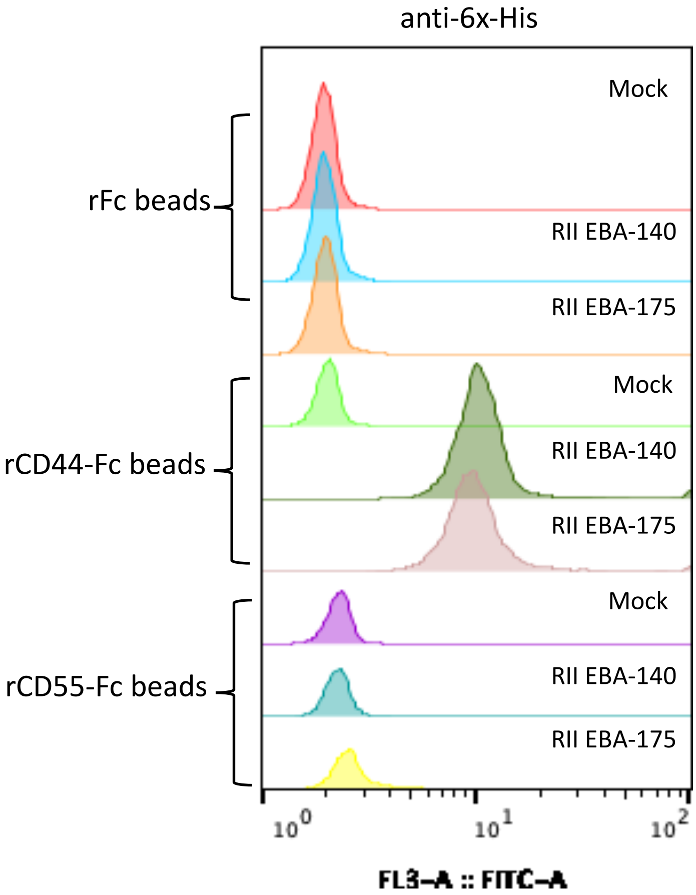
anti-6x-His
rFc beads
rCD44-Fc beads
rCD55-Fc beads
Mock
RII EBA-140
RII EBA-175
Mock
RII EBA-140
RII EBA-175
Mock
RII EBA-140
RII EBA-175
$10^0$
$10^1$
$10^2$
FL3–A :: FITC–A

Figure S2

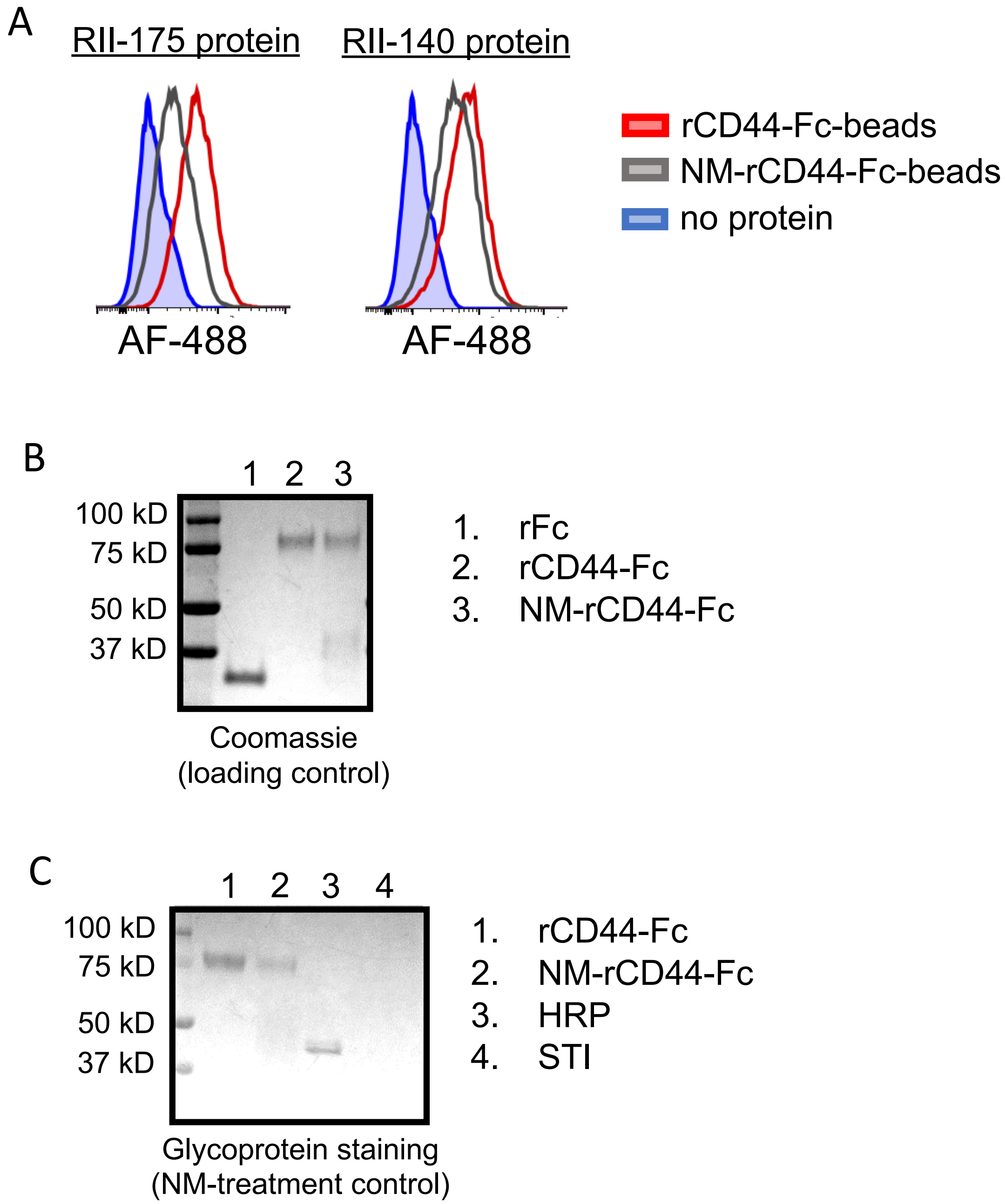

A
RII-175 protein
RII-140 protein
AF-488
AF-488
rCD44-Fc-beads
NM-rCD44-Fc-beads
no protein
B
1 2 3
100 kD
75 kD
50 kD
37 kD
1. rFc
2. rCD44-Fc
3. NM-rCD44-Fc
Coomassie
(loading control)
C
1 2 3 4
100 kD
75 kD
50 kD
37 kD
1. rCD44-Fc
2. NM-rCD44-Fc
3. HRP
4. STI
Glycoprotein staining
(NM-treatment control)

Figure S3

A

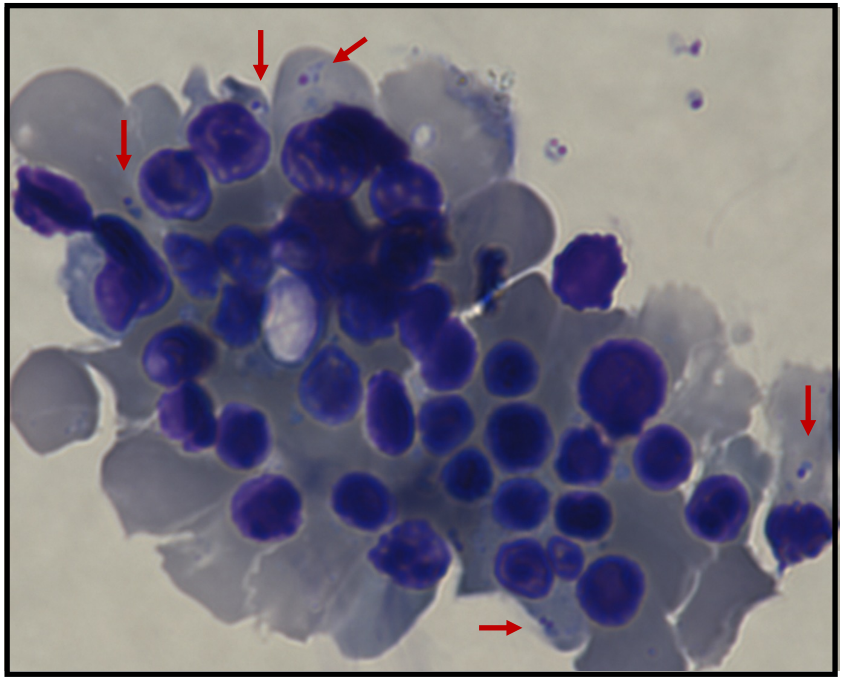

B

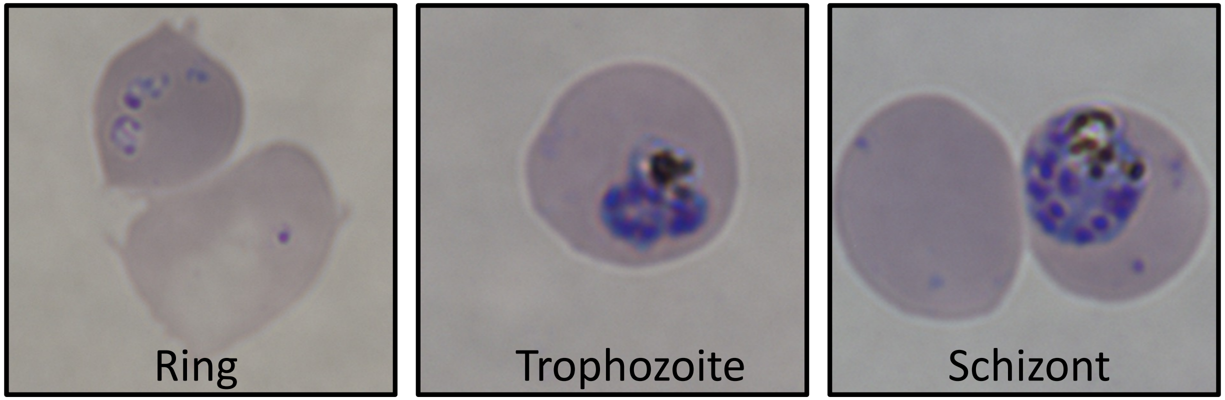

### Figure S4

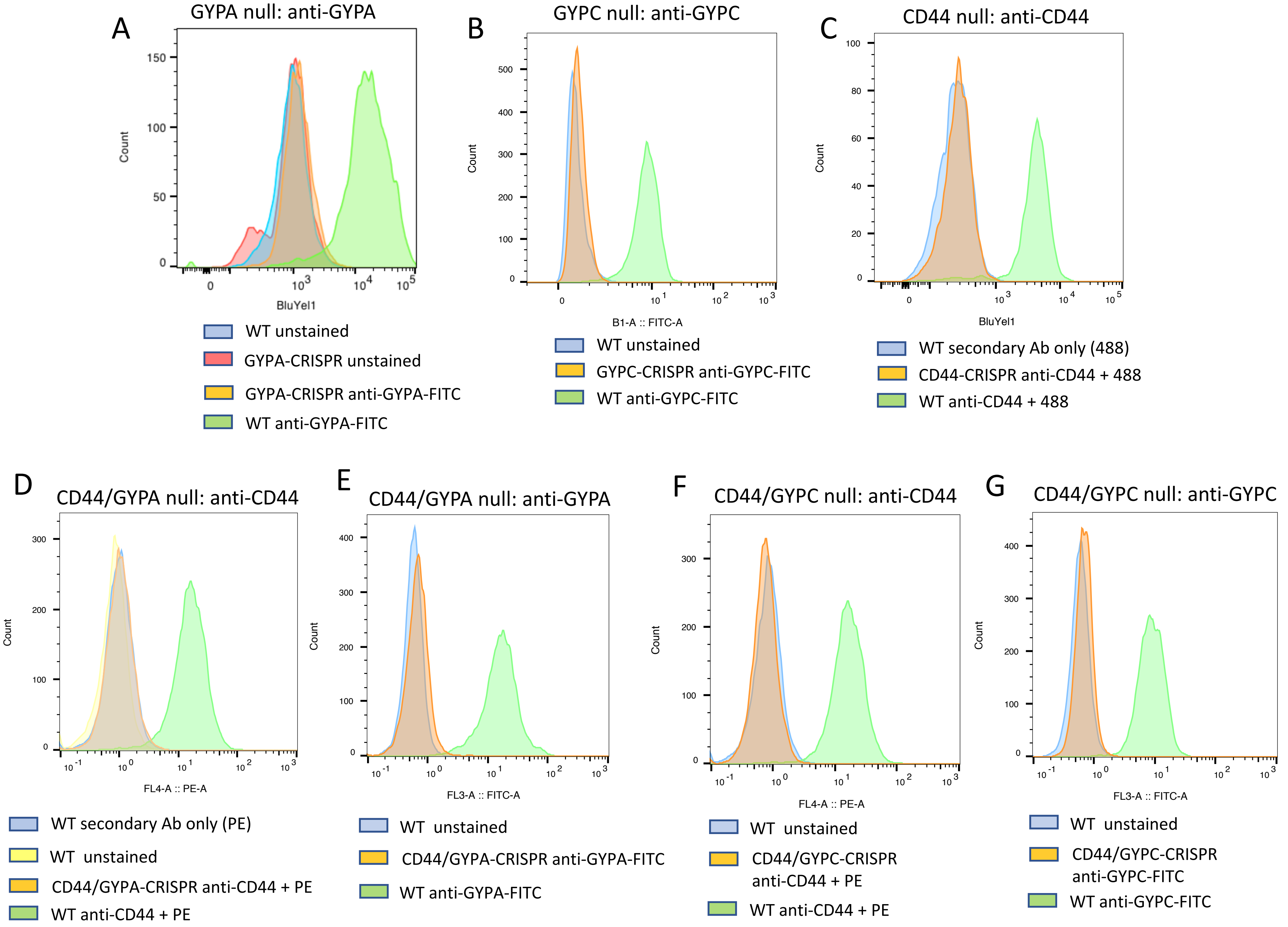
A
GYPA null: anti-GYPA
Count
BluYel1
WT unstained
GYPA-CRISPR unstained
GYPA-CRISPR anti-GYPA-FITC
WT anti-GYPA-FITC
B
GYPC null: anti-GYPC
Count
B1-A :: FITC-A
WT unstained
GYPC-CRISPR anti-GYPC-FITC
WT anti-GYPC-FITC
C
CD44 null: anti-CD44
Count
BluYel1
WT secondary Ab only (488)
CD44-CRISPR anti-CD44 + 488
WT anti-CD44 + 488
D
CD44/GYPA null: anti-CD44
Count
FL4-A :: PE-A
WT secondary Ab only (PE)
WT unstained
CD44/GYPA-CRISPR anti-CD44 + PE
WT anti-CD44 + PE
E
CD44/GYPA null: anti-GYPA
Count
FL3-A :: FITC-A
WT unstained
CD44/GYPA-CRISPR anti-GYPA-FITC
WT anti-GYPA-FITC
F
CD44/GYPC null: anti-CD44
Count
FL4-A :: PE-A
WT unstained
CD44/GYPC-CRISPR anti-CD44 + PE
WT anti-CD44 + PE
G
CD44/GYPC null: anti-GYPC
Count
FL3-A :: FITC-A
WT unstained
CD44/GYPC-CRISPR anti-GYPC-FITC
WT anti-GYPC-FITC

### Figure S5

A

Total protein | Phospho-tag | Overlay

WT Mock (gel 5)

WT + EBA 140 (gel 6)

ΔCD44 Mock (gel 7)

ΔCD44 + EBA 140 (gel 8)

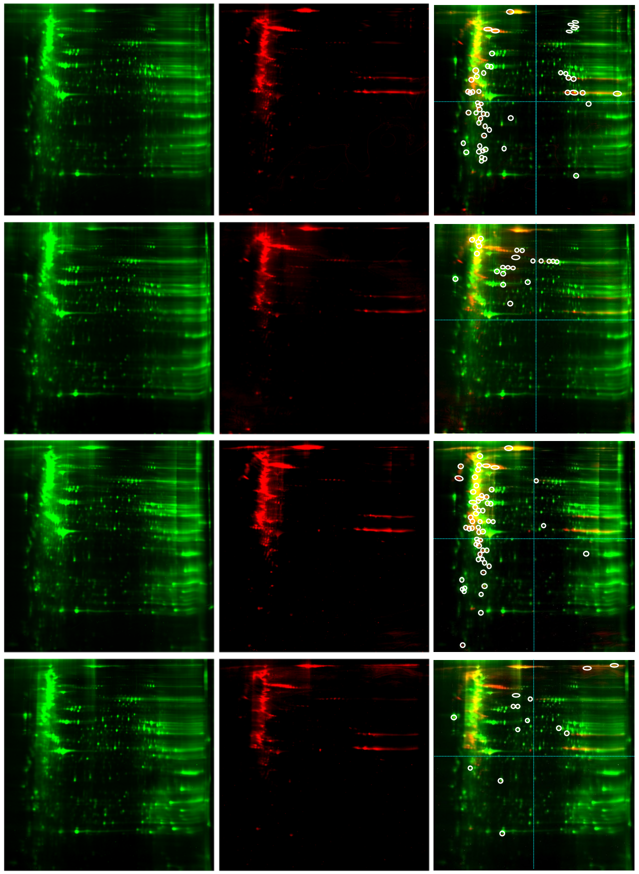

B

WT + EBA 140 (gel 6 overlay) | ΔCD44 + EBA 140 (gel 8 overlay)

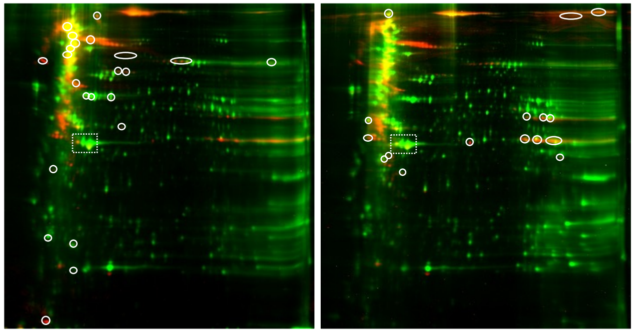

C

WT Mock (gel 5) | WT + EBA 140 (gel 6)

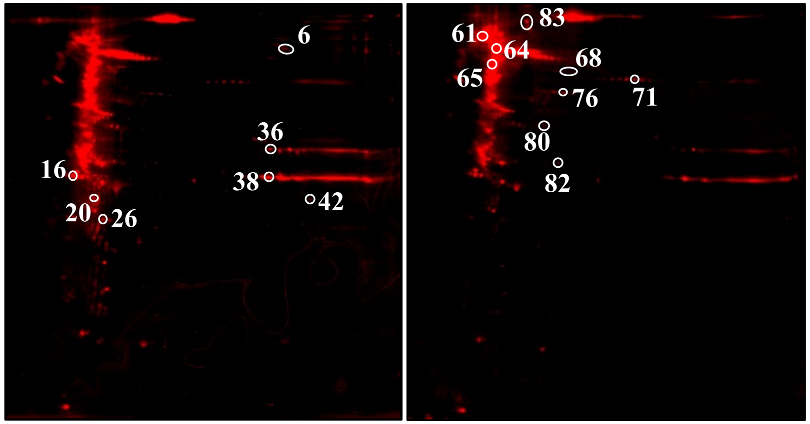


D

| Spot | | Name | protein score CI % | Total Ion CI% |
|---|---|---|---|---|
| 6 | ANK1 | Ankyrin-1 | 100 | 100 |
| | ACLY | ATP-citrate synthase | 100 | 100 |
| 16 | ADRM1 | Proteasomal ubiquitin receptor | 100 | 100 |
| | SPTB1 | Spectrin beta chain | 100 | 100 |
| | SPTA1 | Spectrin alpha chain | 99.744 | 99.997 |
| | RINI | Ribonuclease inhibitor | 99.262 | 99.95 |
| 20 | SPTB1 | Spectrin beta chain | 100 | 100 |
| 26 | IF2A | Eukaryotic translation initiation factor 2 subunit 1 | 100 | 100 |
| 36 | ATPA | ATP synthase subunit alpha | 100 | 100 |
| | 41 | Protein 4.1 | 99.956 | 99.969 |
| 38 | OAT | Ornithine aminotransferase | 100 | 100 |
| | PUR6 | Multifunctional protein ADE2 | 100 | 100 |
| 42 | GNA13 | Guanine nucleotide-binding protein subunit alpha-13 | 100 | 100 |
| | ALDOA | Fructose-bisphosphate aldolase A | 100 | 100 |
| 61 | SPTB1 | Spectrin beta chain | 100 | 100 |
| | SPTA1 | Spectrin alpha chain | 100 | 100 |
| 64 | SPTB1 | Spectrin beta chain | 100 | 100 |
| | SPTA1 | Spectrin alpha chain | 99.997 | 99.491 |
| 65 | SPTA1 | Spectrin alpha chain | 100 | 100 |
| | SPTB1 | Spectrin beta chain | 100 | 100 |
| 68 | MIC60 | MICOS complex subunit MIC60 | 99.613 | 99.829 |
| 71 | 41 | Protein 4.1 | 100 | 100 |
| | E41L1 | Band 4.1-like protein 1 | 100 | 100 |
| | E41L3 | Band 4.1-like protein 3 | 100 | 100 |
| 76 | GRP75 | Stress-70 protein, mitochondrial | 100 | 100 |
| | K2C1 | Keratin, type II cytoskeletal 1 | 99.992 | 100 |
| 80 | TCPQ | T-complex protein 1 subunit theta | 100 | 100 |
| 82 | IF2B | Eukaryotic translation initiation factor 2 subunit 2 | 100 | 100 |
| 83 | B3AT | Band 3 anion transport protein | 100 | 100 |
| | ANK1 | Ankyrin-1 | 100 | 100 |

**Supplementary Figures**

**Figure S1. Additional evidence showing *P. falciparum* invasion ligands EBA-175 and EBA-140 interact directly with CD44.** rCD44-Fc or rCD55-Fc made in human HEK293 cells were bound to protein A beads and incubated with RII of EBA-175-His or RII of EBA-140-His. rFC-coated beads were used as a control. Mock conditions were incubated in PBS instead of EBA protein. Binding of the EBA proteins to the protein-coated beads was quantified by flow cytometry using an anti-6x-His mouse monoclonal antibody and goat anti-mouse-FITC secondary antibody.

**Figure S2. The interactions between CD44 and EBA-175 or EBA-140 are partially dependent on sialic acid.** A) rCD44-Fc made in a mouse myeloma cell line (R&D) was treated with neuraminidase (NM) to remove sialic acid. Beads coated with NM-treated rCD44-Fc or untreated rCD44-Fc were used for *in vitro* binding to RII EBA-175-His and RII EBA-140-His. Uncoated beads (no protein) were used as a negative control. Binding of the EBA proteins to the beads was quantified by flow cytometry using an anti-6x-His mouse monoclonal antibody.
B) Coomassie gel showing similar concentrations of rCD44-Fc and NM-treated rCD44-Fc (NM-rCD44-Fc). C) Gel analysis of glycoproteins shows reduced staining of NM-rCD44-Fc relative to rCD44-Fc. HRP and STI are positive and negative controls provided in the glycoprotein staining kit, respectively (see methods).

**Figure S3. *P. falciparum* can invade and develop in cRBCs derived from HUDEP-2 cells.** A) HUDEP-2 cells were induced to differentiate down the erythroid lineage to orthochromatic erythroblasts, and incubated with *P. falciparum* schizonts. After ~18 hours, ring-stage parasites were detected in nucleated cRBCs on cytospin slides stained with May-Grünwald and Giemsa, as indicated by red arrows. B) *P. falciparum* can progress through its asexual cycle and develop into schizonts in enucleated HUDEP-2 cRBCs.

**Figure S4. Validation of HUDEP-2 mutant clones by flow cytometry.** A) GYPA-null HUDEP-2 or WT HUDEP-2 stained with anti-GYPA. B) GYPC-null HUDEP-2 or WT HUDEP-2 stained with anti-GYPC. C) CD44-null HUDEP-2 or WT HUDEP-2 stained with BRIC 222.

D) GYPA/CD44-null HUDEP-2 or WT HUDEP-2 stained with BRIC 222. E) GYPA/CD44-null HUDEP-2 or WT HUDEP-2 stained with anti-GYPA. F) GYPC/CD44-null HUDEP-2 or WT HUDEP-2 stained with BRIC 222. G) GYPC/CD44-null HUDEP-2 or WT HUDEP-2 stained with anti-GYPC.

**Figure S5. 2D-DIGE gel analysis of ghost lysates from WT or CD44-null cRBCs stimulated with EBA-140.** A) WT or CD44-null cRBCs were stimulated with mock vs. RII EBA-140. Each sample was labelled separately with dyes for total protein and fluorescent phospho-tag, which were overlaid on the same 2D-DIGE gel. DeCyder software was used to quantify phosphorylated spots in each gel, and to identify spots with differential phosphorylation between Mock and EBA-140 stimulation in each genetic background (white circles indicate increased phosphorylation). B) Gel 6 and 8 overlays were manually aligned to identify proteins with differential phosphorylation between WT and CD44-null cRBCs. White circles indicate spots with increased phosphorylation relative to the other genetic background. White box denotes normalization control. C) Spots picks for identification by mass spectrometry, based on intersection of spots with altered phosphorylation upon EBA-140 stimulation (gels 5 & 6) with spots in which phosphorylation depends on CD44 (gels 6 & 8). D) 16 spots picked for protein identification by mass spectrometry at Applied Biomics, Inc. The Protein Score C.I. % is the confidence of the protein ID calculated from MS data. The Total Ion C.I. % is the confidence of the protein ID calculated from MS/MS data. Scores above 95% are significant for both.
